# Exploring Synergies in Brain-Machine Interfaces: Compression vs. Performance

**DOI:** 10.1101/2025.02.03.636273

**Authors:** Luis H. Cubillos, Madison M. Kelberman, Matthew J. Mender, Aren Hite, Hisham Temmar, Matthew Willsey, Nishant Ganesh Kumar, Theodore A. Kung, Parag G. Patil, Cynthia Chestek, Chandramouli Krishnan

**Author notes:** These authors contributed equally (Luis H. Cubillos, Madison M. Kelberman). Indicates co-corresponding authors (Cynthia Chestek, Chandramouli Krishnan). Corresponding Author: Chandramouli Krishnan, PT, PhD University of Michigan 325 E Eisenhower Parkway, Suite 3013 Ann Arbor, MI – 48108.

## Abstract

Individuals with severe neurological injuries often rely on assistive technologies, but current methods have limitations in accurately decoding multi-degree-of-freedom (DoF) movements. Intracortical brain-machine interfaces (iBMIs) use neural signals to provide a more natural control method, but currently struggle with higher-DoF movements—something the brain handles effortlessly. It has been theorized that the brain simplifies high-DoF movement through muscle synergies, which link multiple muscles to function as a single unit. These synergies have been studied using dimensionality reduction techniques like principal component analysis (PCA), non-negative matrix factorization (NMF), and demixed PCA (dPCA) and successfully used to reduce noise and improve offline decoder stability in non-invasive applications. However, their effectiveness in improving decoding and generalizability for implanted recordings across varied tasks is unclear. Here, we evaluated if brain and muscle synergies can enhance iBMI performance in non-human primates performing a two-DoF finger task. Specifically, we tested if PCA, dPCA, and NMF could compress and denoise brain and muscle data and improve decoder generalization across tasks. Our results showed that while all methods effectively compressed data with minimal loss in decoding accuracy, none improved performance through denoising. Additionally, none of the methods enhanced generalization across tasks. These findings suggest that while dimensionality reduction can aid data compression, alone it may not reveal the “true” control space needed to improve decoder performance or generalizability. Further research is required to determine whether synergies are the optimal control framework or if alternative approaches are required to enhance decoder robustness in iBMI applications.

**Significance Statement:** Many researchers believe that brain and muscle synergies represent a fundamental control strategy and could enhance brain-machine interface (BMI) decoding performance. These synergies, extracted through dimensionality reduction techniques, are thought to simplify complex neural data, improving the efficiency and accuracy of BMI systems. In our study, we evaluated brain and muscle synergies in a dexterous finger task. We found that while these synergies effectively compressed high-dimensional data, they did not improve performance through denoising or generalize well across different contexts. Instead, the highest performance was achieved when using all available data, suggesting that synergies, although useful for data compression, may not provide the “true” control space needed to enhance decoder robustness or adaptability in implanted BMI systems.

## Introduction

Individuals with serious neurological injuries (e.g., spinal cord injury, severe stroke) often require assistive technologies to perform basic activities of daily living (Maresova et al., 2020).

Intracortical brain-machine interfaces (iBMIs) have appeared as a promising technology for these individuals (Lobel and Lee, 2014). iBMIs offer a natural way of controlling external devices by directly decoding brain signals into behavioral commands, and have been successfully applied in various settings, including controlling robotic arms (Hochberg et al., 2012; Flesher et al., 2021) and decoding live speech (Willett et al., 2021; Card et al., 2024). Although impressive, iBMIs have enabled simultaneous control of only a few DoFs (Deo et al., 2024; Shah et al., 2024; Willsey et al., 2024)—whereas the human brain can control hundreds of muscles at the same time for fluid motion.

One hypothesis for the brain’s efficiency in managing multiple degrees of freedom is that the nervous system simplifies this complexity by functionally grouping muscles, called muscle synergies, to act as a single unit (Holdefer and Miller, 2002; d’Avella et al., 2003; Tresch and Jarc, 2009; Overduin et al., 2015). Although the anatomical basis of muscle synergies is debated (Tresch and Jarc, 2009; Kutch and Valero-Cuevas, 2012; Ranganathan and Krishnan, 2012; Ranganathan et al., 2016), the concept has proven practically useful for understanding the neural control of movement. To investigate these muscle synergies, researchers have used dimensionality reduction (DR) techniques (e.g., principal component analysis [PCA], non-negative matrix factorization [NMF]) on muscle data recorded across various tasks, including walking (Chvatal and Ting, 2012), reaching (d’Avella et al., 2006), and grasping (Overduin et al., 2008). Similarly, these findings have been extended to neural populations in the brain (i.e., brain synergies, usually referred to as neural modes that form a neural manifold in the literature (Gallego et al., 2018)), suggesting that DR might reveal the underlying signals of neural circuits during different tasks (Gallego et al., 2018; Degenhart et al., 2020). If muscle and brain synergies are indeed a fundamental component of the brain’s control strategy, then they could serve as an optimal control space for brain-machine interfaces, leveraging the brain’s inherent structure to facilitate effective control. However, this remains an open question: if synergies are more than an abstraction, they should ideally provide a perfect framework for iBMI control, allowing us to capitalize on the brain’s natural control strategies for improving decoding performance.

Brain and muscle synergies obtained through DR have already proven to be useful in many applications. They have reduced noise in noninvasive recordings (both artifact-induced and electrical noise (Winkler et al., 2011; Damon et al., 2013; Islam et al., 2021; Costa-García et al., 2023)), enhanced iBMI decoder stability across sessions (Degenhart et al., 2020; Karpowicz et al., 2022), and compressed high-dimensional physiological data (Cozza et al., 2020). However, it is unclear whether the denoising benefits would transfer to implanted recordings, whether dimensionality reduction methods can help adapt to new task contexts beyond their initial training, and whether their compression benefits vary between methods and physiological origin. Addressing these issues and comparing different methods could enhance the functionality and clinical translatability of BMI and neuroprosthetic interfaces.

In this study, we examined the potential of brain and muscle synergies extracted from three popular dimensionality reduction techniques to enhance intracortical BMI performance in predicting finger kinematics or forearm muscle activity in non-human primates (NHPs). We systematically assessed their ability to compress the data while maintaining performance, denoise the data by increasing the specificity of the recording, and facilitate generalization across different task contexts. We hypothesized that we could greatly reduce the dimensionality of brain and muscle data without losing performance and that removing some of the lower-variance dimensions would help remove noise and improve iBMI performance. Overall, we found that dimensionality reduction methods were useful for data compression, with dPCA showing the best performance, but did not show denoising or generalization benefits.

## Materials and Methods

### Experimental Design

#### Task and Data Acquisition

We implanted two non-human primates (NHPs), monkeys N and W, with Utah microelectrode arrays in the pre-central gyrus at the arcuate sulcus, which is an anatomic landmark for the ‘hand area’ of motor cortex in NHPs. Monkey N was implanted with two 64-channel arrays, while Monkey W received a 96-channel array, and both recordings were limited to 96 channels. Additionally, we implanted Monkey N with eight intramuscular bipolar electromyographic (EMG) electrodes in the forearm, targeting muscles involved in finger and wrist movements (Flexor Carpi Radialis (FCR), Flexor Digitorum Profundus (three electrodes, denoted by FDP-distal [FDPd], proximal [FDPp], and FDP), Flexor Carpi Ulnaris (FCU, excluded due to disconnection from target), Extensor Carpi Radialis Brevis (ECRB), Extensor Indicis Propius (EIP), and Extensor Digitorum Communis (EDC); Figure 1A).

**Figure 1:**
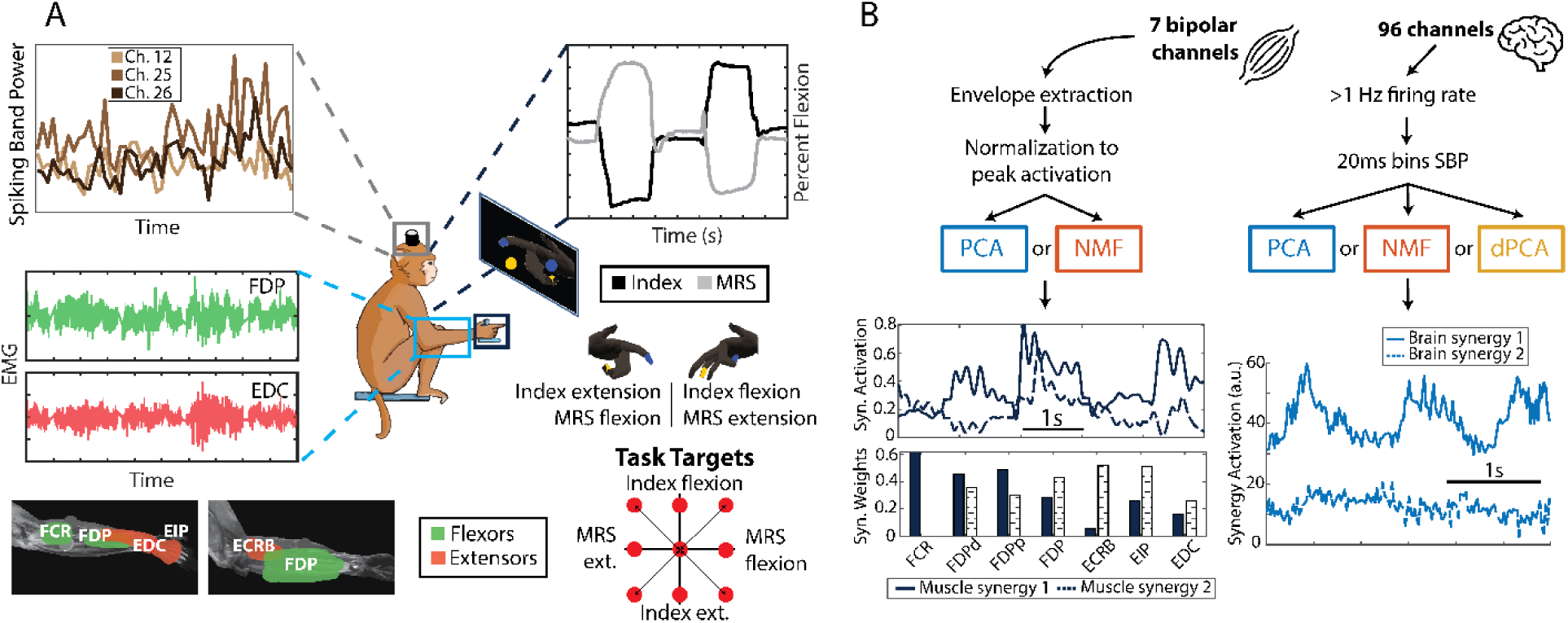
Monkey task and data processing pipeline. (A) Monkeys N and W were trained to do a 2-DoF dexterous finger task, in which they had to move the index and the middle-ring-small (MRS) fingers independently to targets shown on the screen (center diagram). While they did the task, we recorded finger position and velocity (top right plot), spiking band power of the brain channels (top left plot) and, for Monkey N, muscle activity (bottom left plot). (B) Diagram showing the data processing pipeline for the muscle (left) and brain (right) data. The 7 bipolar muscle channels are processed to extract envelopes and then normalized to the peak activation from each day. Then, we applied PCA and NMF, resulting in a set of muscle synergy activations (top plot) and synergy weights (bottom plot). The 96 brain channels are filtered to extract only those with at least 1Hz firing rate on average, and then transformed using the SBP in 20ms bins. PCA, NMF, and dPCA are then applied, which results in a set of brain synergy activations. **Abbreviations**: MRS = middle-ring-small fingers; EMG = electromyography; PCA = principal component analysis; NMF = non-negative matrix factorization; dPCA = demixed PCA; SBP = spiking band power; FDP = flexor digitorum profundus; EDC = extensor digitorum communis; FCR = flexor carpi radialis; ECRB = extensor carpi radialis brevis; EIP = extensor indicis propius.

We trained the monkeys to perform a 2-degree-of-freedom (2-DoF) dexterous finger task. This task required them to flex and extend their index and middle-ring-small (MRS) fingers independently while targets were shown in colors over a virtual hand on a screen in front of the monkeys (screen next to monkey in Figure 1A). Targets were presented in a center-out manner: every two trials, the targets appeared on the center, which corresponds to a resting position between flexion and extension. Successful trial completion required that the monkey maintain its finger positions within the targets for a preset hold time of 750 ms. At the end of each trial, monkeys were rewarded with fruit juice by using a juicer machine that turned on a pump for 100 milliseconds (juicer artifact in Figure 2). Finger flexion was measured using a custom-designed manipulandum equipped with daily-calibrated bend sensors, as described previously (Nason et al., 2021).

**Figure 2:**
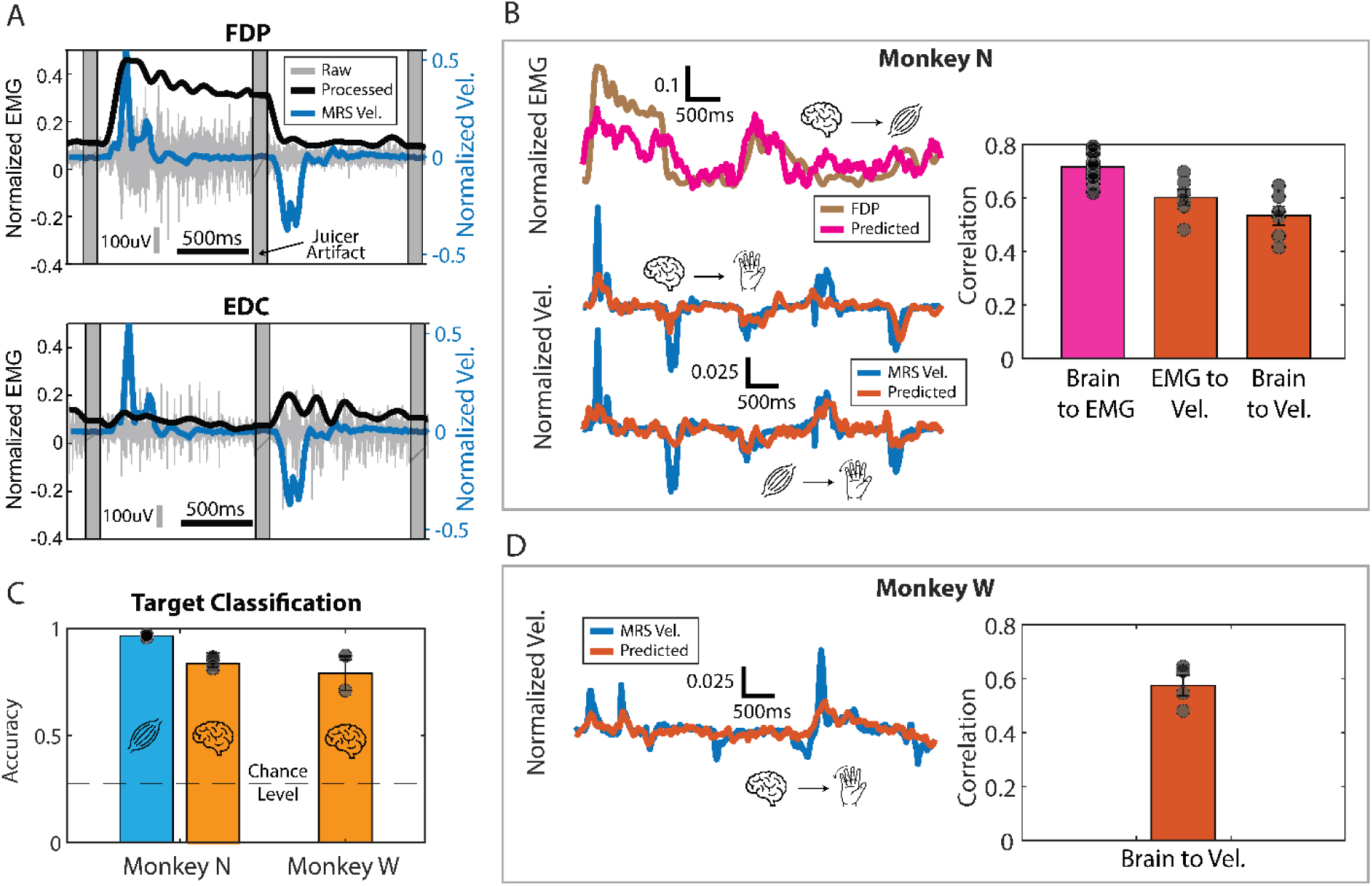
Assessment of the quality of physiological signals. (A) Monkey N’s raw and processed EMG, together with MRS finger velocity for two example trials. Gray vertical bars represent the removed juicer artifact (see Methods) at the beginning of each trial. Top plot shows FDP, a muscle targeting MRS flexion, and the bottom shows EDC, a muscle targeting MRS extension. Vertical gray line shows scale of 100uV for the raw EMG and horizontal black line shows scale for 500ms. (B) Example traces of Monkey N’s processed FDP EMG predicted from brain activity (top) and finger velocity predicted from brain activity (middle) and processed EMG (bottom). Bar plot on the right shows aggregated correlations corresponding to three days of Monkey N data. Black dots in the left bar represent the predictions for each muscle channel (seven dots for each day). Bar plots in middle and right bar represent the velocity predictions for each finger group (two dots for each day). (C) Accuracy of classifying targets using EMG (muscle symbol) and brain activity (brain symbol) for both monkeys across days. Segmented line shows chance level. Aggregated results correspond to three days of Monkey N and two days of Monkey W. (D) Same as in B but for Monkey W. Trace shows example of MRS velocity and the prediction when using the brain data. Bar plot shows aggregated results across 2 days of Monkey W data. **Abbreviations:** FDP = flexor digitorum profundus; EDC = extensor digitorum communis; MRS = middle-ring-small fingers; Vel. = velocity; EMG = electromyography.

Additionally, monkeys performed three variations of the task: one in which the wrist was flexed by 23 degrees (“wrist context”), another in which a spring to resist flexion was added (“spring context”), and a final one that included both changes (“spring+wrist context”). These context changes allowed us to explore changes in brain and muscle activity due to the variations in task while keeping the finger kinematics the same, as shown previously (Mender et al., 2023).

Data acquisition included real-time brain and finger movement data collection via xPC Target (Mathworks), ensuring millisecond precision. These data were later synchronized with EMG data. The 96 brain channels were sampled at 30kHz and bandpass filtered from .3-7500 Hz. Threshold crossings were detected by setting a threshold of −4.5 times the signal root-mean-square (RMS) during an initial calibration recording. For brain data analyses, only channels with at least one threshold crossing per second on average across the recording session were used. For each of these channels, we extracted the spiking band power (power in the 300-1000 Hz band, SBP, (Nason et al., 2020)) feature, known for its low power consumption while maintaining decoding performance. The SBP and finger movements were then averaged into 20ms bins (Figure 1B, right). On the other hand, the EMG data processing pipeline involved downsampling the original signal to 2000Hz using an 800Hz anti-aliasing filter, followed by a band-pass filter (100 to 500Hz), signal rectification, and a final low-pass filter at 6Hz to extract the envelopes (“Envelope Extraction” in Figure 1B, left) (Gallego et al., 2018; Mender et al., 2023). The processed EMG was then normalized to the peak activation of each day and the resulting signal is referred to as the muscle activity throughout the paper. The binned SBP and kinematics, as well as the muscle activity were used for all analyses, as opposed to the raw data for each.

#### Dimensionality Reduction Methods

We tested the application of different dimensionality reduction methods to the brain and muscle data. These methods work by taking an N-dimensional time series (such as the spiking band power of 96 channels throughout an experiment; N = 96 in that case) and extracting M (M <= N) synergies. These synergies can usually be described with two components: the synergy activations and the synergy weights (Figure 1B, bottom). The synergy activations are M-dimensional time series representing the corresponding synergy activity. The synergy weights, on the other hand, represent the relative scaling of each of the original N dimensions for the corresponding synergy. In the neural population literature, the synergy weights are called neural modes, which form a neural manifold, and the synergy activations are called the neural trajectories of those modes in the neural manifold.

We applied three dimensionality reduction methods to the data: PCA (Abdi and Williams, 2010), NMF (Rabbi et al., 2020), and dPCA (Kobak et al., 2016). All three were applied to brain activity, but only PCA and NMF were applied to the muscle activity, as dPCA’s main benefit is extracting movement-relevant components from brain data (Kobak et al., 2016). All three methods identify low-dimensional components that can reconstruct the original data as best as possible, but they differ in the specific loss functions and their restrictions. First, PCA defines a decoder/encoder matrix D and tries to minimize the reconstruction error of the data *X* in the following form:

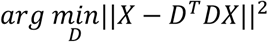

In this study, we denominated *DX* as the synergy activations (a.k.a. scores) and *D*^*T*^ as the synergy weights (a.k.a. loadings). The *D* matrix can be solved analytically, making PCA decomposition a fast process. NMF, on the other hand, imposes a non-negative constraint on the weights (*W*) and the activation (*H*), and minimizes the reconstruction error of the data *X* in the following form:

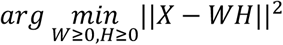

NMF does not have an analytical solution; rather, numerical methods are needed to solve for *W* and *H*, making it a more demanding computation than for PCA. NMF is frequently used because it may better represent physiological signals due to its non-negative constraint (Rabbi et al., 2020). Finally, dPCA is similar to PCA, but it is supervised (i.e., has access to the behavior data for training) and also relaxes the restriction that the encoder and decoder transformations must be the same, which allows for selection of principal components in the brain activity most related to pre-specified task parameters. Like PCA, the dPCA matrices can also be solved analytically, using a similar objective function which accounts of task parameters, seen below:

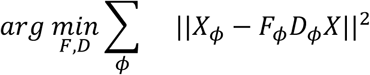

Where *ϕ* represents the task parameter (i.e., target position or time), *F*_*ϕ*_ is the encoder matrix for that parameter, *D*_*ϕ*_ is the decoder matrix for that parameter, and *X*_*ϕ*_ is the trial-averaged data separated with regards to that parameter. In this study, we used two task parameters to separate the data: target position (nine possible values given by all combinations of flex, extend, and rest for each finger group; see Task Targets in Figure 1A) and time (i.e., a condition-independent parameter), both of which are common choices when using dPCA (Kobak et al., 2016). The synergy activation for dPCA was determined by computing *D*_*ϕ*_*X*, with *D*_*ϕ*_being the decoder matrix related to the target location, and *X* the non-parameter-separated data.

### Statistical Analyses

#### Signal Quality Assessment

To assess the quality of the recorded physiological signals, we determined whether they were related to finger movements during the task. This tested the predictive ability of brain activity to estimate muscle activity and finger kinematics and of muscle activity to predict finger kinematics. For three days of 500 trials each with Monkey N, we trained three ridge-regression-with-history models (Collinger et al., 2013): one to predict muscle activity from brain activity, another to predict finger velocity from brain activity, and the last to predict finger velocity from muscle activity. For two days of 500 trials each with Monkey W, since only brain and kinematics data were available, we trained a single model predicting finger velocity from brain activity. All models had a total of 10 bins of history, equivalent to 200ms. Additionally, to study movement discrimination ability in the recorded signals, we trained linear classifiers (support vector machines with a linear kernel, *fitcecoc* in Matlab) that distinguished between nine different task targets (all combinations between flex, extend, and rest for each finger group; Figure 3A). The performance of the ridge regression models was measured with the Pearson correlation coefficient between the ground truth and the prediction, and the classifier’s performance with the average accuracy across targets.

**Figure 3:**
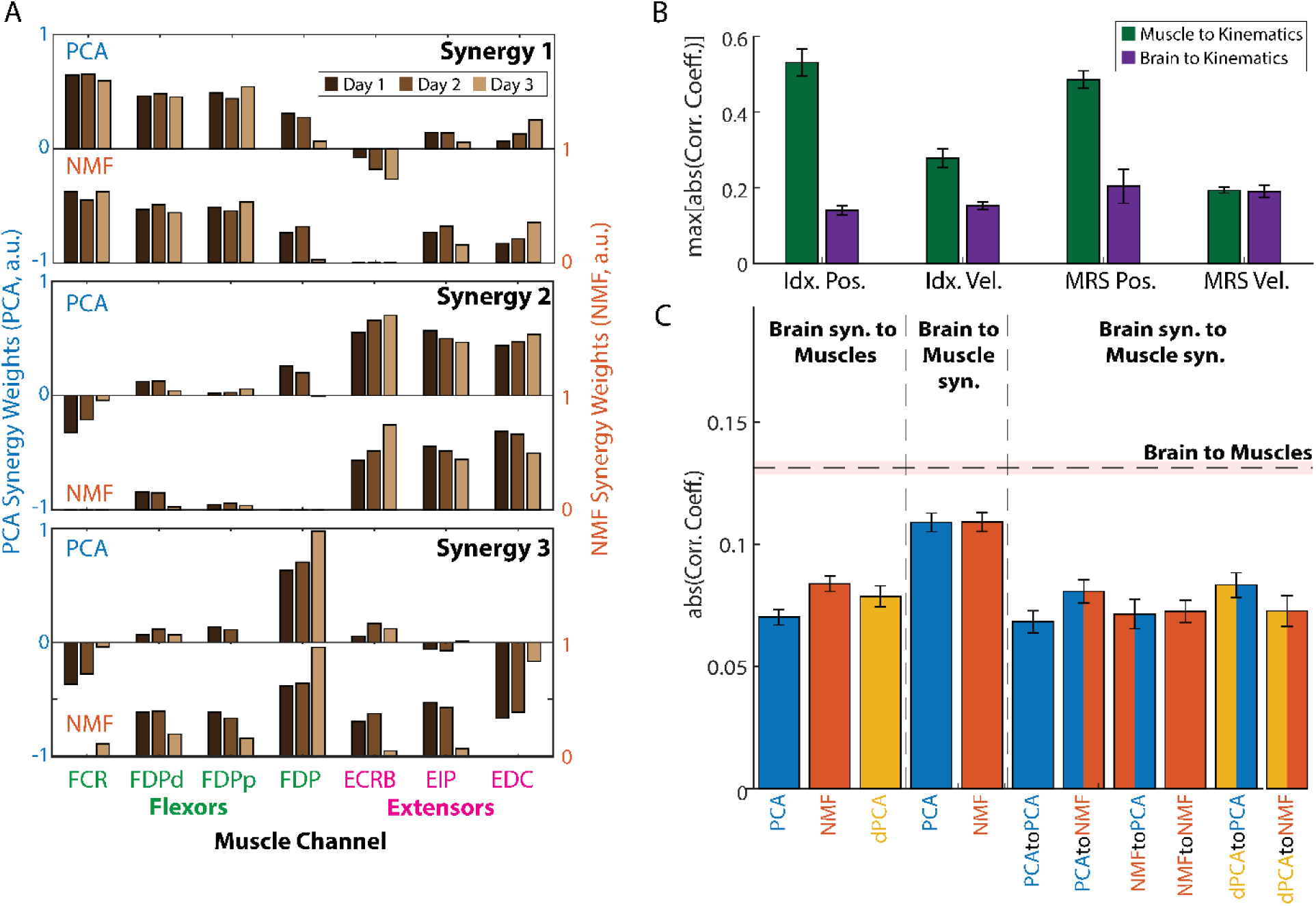
Analysis of brain and muscle synergy relationships for monkey N. (A) Muscle synergy weights extracted using PCA (top bar plot in each window) and NMF (bottom bar plot) across days. Synergy weights were consistent across days and methods and revealed a structure closely related to the task. (B) Barplot showing the average across days of the maximum absolute correlation coefficient between all muscle channels and all brain channels with the four kinematic outputs: index and MRS position, index and MRS velocity. The brain channels had a weaker direct relationship with the kinematic outputs, which hinders an interpretation of the brain synergies similar to that shown in A. (C) Average of the absolute correlation coefficient between all possible pairs of brain and muscle synergy activations, broken down by the method(s) used. The black segmented line shows the average of the absolute correlation coefficient between all pairs of brain and muscle channels, while the shading represents the standard error of the mean (SEM). All error bars represent the SEM as well. **Abbreviations**: PCA = principal component analysis; NMF = non-negative matrix factorization; FCR = flexor carpi radialis; FDP = flexor digitorum profundus; FDPd = FDP distal; FDPp = FDP proximal; ECRB = extensor carpi radialis brevis; EIP = extensor indicis propius; EDC = extensor digitorum communis; Idx = index finger; MRS = middle-ring-small fingers; Pos. = position; Vel. = velocity; Syn. = synergy; Corr. Coef. = Pearson correlation coefficient; a.u. = arbitrary units.

#### Relation Between Synergies and Movement

After assessing the quality of the signals, we applied dimensionality reduction methods to the brain and muscle data, and explored whether those tools could advance our understanding of the relationship between brain and muscle synergies with movement. Unlike muscle data, which has a more direct and well-understood link to movement (e.g., FDPp is involved in the flexion of the index finger), brain signals can represent complex neural activity that doesn’t always correspond in a straightforward way to intended actions, adding a layer of ambiguity and complexity in decoding. We quantified this brain and muscle interpretability hypothesis by computing the Pearson correlation coefficients between each muscle and brain channel with each kinematic output: index position and velocity and MRS position and velocity. Higher correlations between kinematics and muscle activity, than between kinematics and brain activity, may mean a more direct relationship between muscle activity and kinematics.

Then, we studied the relationship between brain and muscle and their respective synergies. We first determined the number of brain and muscle synergies needed to achieve at least 90% VAF for PCA (Hug et al., 2010; Frère and Hug, 2012; Steele et al., 2015; Turpin et al., 2021). Given a set of muscle and brain synergies for each method, we computed the Pearson correlation coefficient between every combination of brain channels and synergies with every combination of muscle channels and synergies. For example, for brain channel 1, we computed the correlation with all seven muscle channels, all PCA muscle synergy activations, and all NMF muscle synergy activations. If the correlations with the muscle channels are higher overall than with the muscle synergies, it may suggest that muscle activity is better represented in brain channel 1 than muscle synergies. To determine the significance threshold for correlations, we ran 1000 permutations of the signals for each pair of the computed correlations. We computed the thresholds with an alpha level of 5% and corrected for the number of comparisons using the Bonferroni correction. Finally, we organized the correlations into four groups: (1) brain channels vs. muscle channels, (2) brain channels vs. muscle synergies, (3) brain synergies vs. muscle channels, and (4) brain synergies vs. muscle synergies. We compared the overall correlations between groups using the *cocor* package in R.

#### Compression Experiment

Next, we tested whether PCA, NMF, and dPCA were useful in compressing brain and muscle features when predicting muscle activity and kinematics. First, we computed the variance accounted for (VAF), as described in (Gallego et al., 2018), for each method with both brain and muscle data, with varying numbers of synergies. Then, we trained ridge regression models that took varying numbers of brain or muscle synergies as input and predicted muscle activity and kinematics (when using brain synergies) or only kinematics (when using muscle synergies). We measured their performance by computing the Pearson correlation coefficient between the predictions and the ground truth. Additionally, we trained ridge regression models using the full high-dimensional brain or muscle data as input and compared their performance in predicting muscle activity and kinematics to that of ridge regressions trained with synergies. Given that the performance of these regressions may not necessarily exhibit a strictly monotonic increase, we determined for each the number of synergies necessary to achieve at least 95% of the correlation given by the full-data-trained model by fitting the following exponential function to the results:

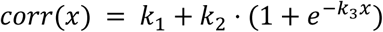

where *x* is the number of synergies used, *corr* represents the Pearson correlation coefficient, and *k*_1_through *k*_3_were fit using gradient-based optimization. After fitting the model, we found the *x* such that *corr(x)* went over 95% of the maximum value (*k*_1_ + *k*_2_). Then, we expressed the compression capabilities of each method for each input as the ratio between the total number of dimensions and this *x*. We tested the compression on three days for Monkey N and two days for Monkey W.

#### Denoising Experiment

Previous research has shown that removing dimensions of brain data (in modalities such as EEG or MEG) can help in decoding by eliminating noise (Winkler et al., 2011; Islam et al., 2021) (a.k.a., denoising). In this case, noise is defined as the electrical activity not related to the behavior of interest (e.g., movement artifacts, power line interference, or neural activity not directly related to behavior). We are interested in studying whether extracting synergies and then using them for decoding can help in denoising our intracortical and intramuscular data. We tested this hypothesis by training multiple models that evaluated muscle activity and kinematics prediction from brain and muscle synergies and compared them to the baselines of using all the available data. If the dimensionality reduction methods can denoise the data, then we would see better predictions coming from a reduced number of synergies than using all the data available. On the other hand, if the methods cannot denoise the data, then removing information, in the form of removing higher-order synergies, will reduce performance. To determine this, we performed the following analysis. We trained eight ridge regressions with different inputs and outputs (Table 1, schematic in Figure 4A). Then, we compared the results of reducing the dimensionality using PCA (brain and muscles), NMF (brain and muscles), and dPCA (only brain), versus just using the full dimensional data. Note that for the models that predict muscle synergy activation rather than muscle activity, an additional layer was added that projects the output into muscle space using the appropriate muscle synergy weights, to compare the predictions across models in muscle space. To measure the performance of these models, we used the Pearson correlation coefficient between the predictions and the ground truth output. We tested denoising on three days for Monkey N, and two days for Monkey W.

**Figure 4:**
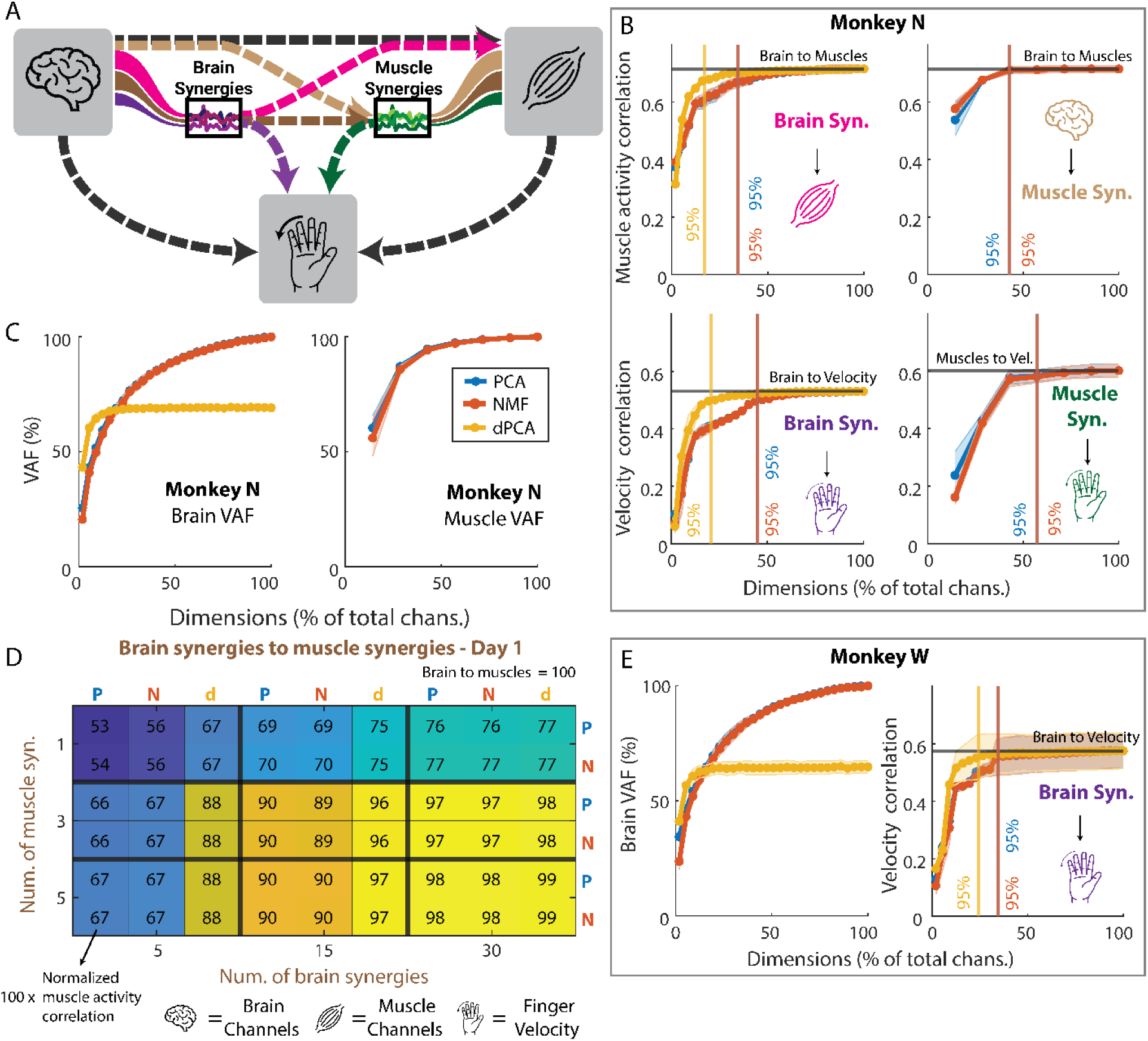
Compression and denoising analyses. (A) Diagram explaining the connections that were tested. Each line represents a different model. For example, the pink line represents a model that was trained with brain synergy activations as the input, and with muscle activity as the output. The three icons represent the brain channels, the muscle activity, and the finger velocity, respectively. The two black lines represent the baselines, shown in black in the rest of the plots. (B, top) Correlation coefficient of the prediction of muscle activity from brain synergies (left) and from the brain channels through muscle synergies (right). The x axis represents what percent of the total dimensions were used to make the predictions. The black lines represent the result of using all brain channels to predict all muscle activity. The colors of the icons are as in A. The vertical lines represent the percent of total dimensions that are needed for each method to achieve 95% of the maximum correlation. The shading for each line represents the standard error of the mean across days. (B, bottom) Correlation coefficient of the prediction of finger velocity from brain synergies (left) and from muscle synergies (right). (C) Average variance accounted for as a function of the percentage of total brain (left) and muscle (right) channels for monkey N. (D) Sample heatmap showing, for day 1, the correlation of predicting muscle activity through brain and muscle synergies, normalized to the result of predicting muscle activity directly from brain channels. A value of 100 would mean that it matches the baseline perfectly. P represents PCA, N represents NMF, and d represents dPCA. (E) Same as B and C but for Monkey W. **Abbreviations**: PCA = principal component analysis; NMF = non-negative matrix factorization; dPCA = demixed PCA; VAF = variance accounted for; Syn. = synergy; Vel. = velocity.

**Table 1:** Ridge regressions for denoising analyses.

| Number | Input | Output |
| --- | --- | --- |
| 1 | Brain activity | Muscle activity |
| 2 | Brain activity | Finger Velocity |
| 3 | Brain activity | Muscle syn. activation |
| 4 | Brain syn. activation | Muscle activity |
| 5 | Brain syn. activation | Finger Velocity |
| 6 | Brain syn. activation | Muscle syn. activation |
| 7 | Muscle activity | Finger Velocity |
| 8 | Muscle syn. activation | Finger Velocity |

#### Generalization Experiment

Finally, previous studies have shown that latent spaces extracted with dimensionality reduction methods are stable across days and tasks (Gallego et al., 2018), and we wanted to test whether they could help for decoding generalizing across task contexts. We had Monkey N perform the finger task in four different contexts (normal, spring, wrist, and spring+wrist) sequentially and then trained ridge regression models to predict finger velocity and muscle activity for each non-normal context. Off-context models refer to those trained using 100% of the data in the normal context and then tested using the last 20% of the data in each non-normal context (Figure 5A).

**Figure 5:**
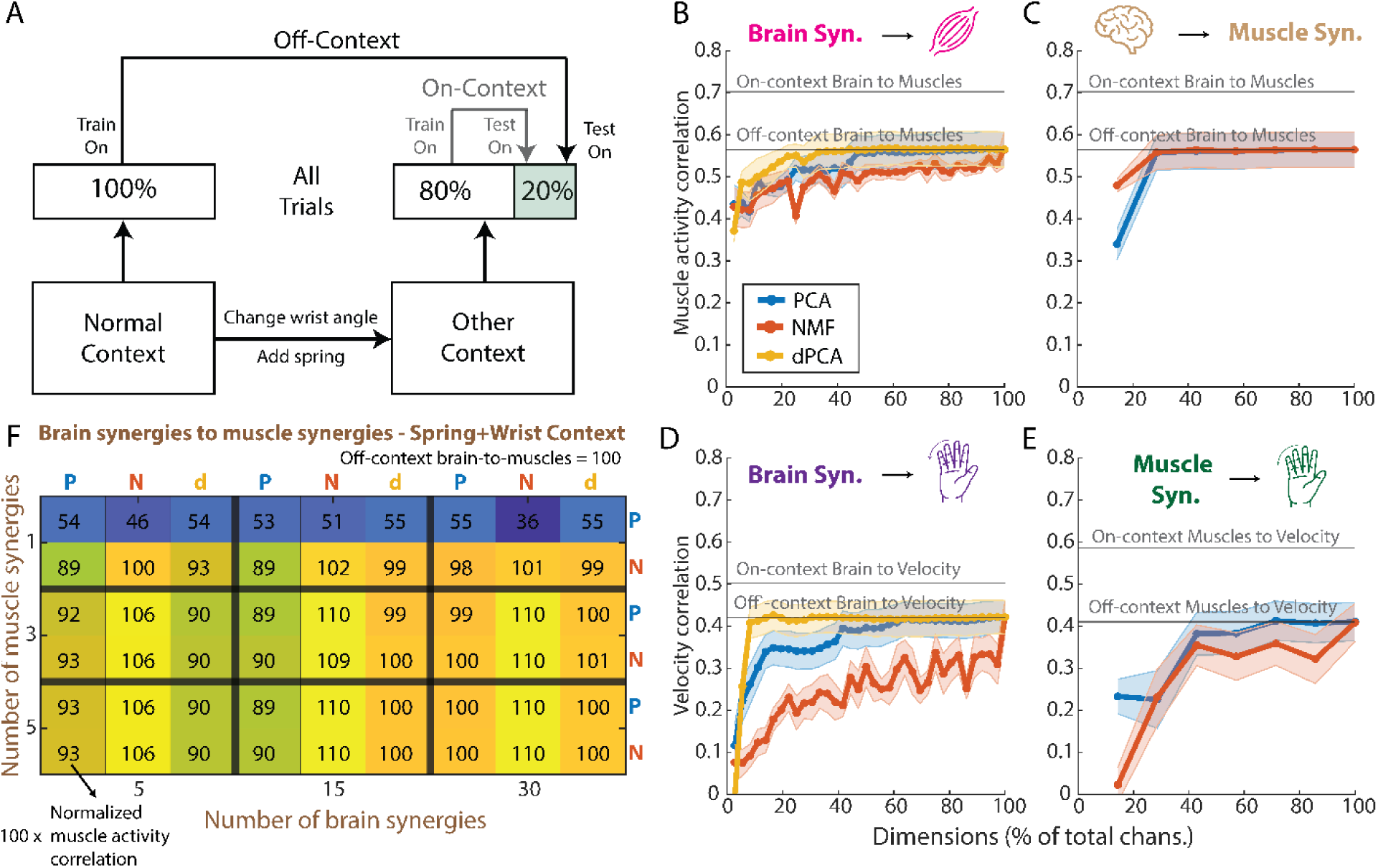
Generalization analyses for Monkey N. (A) Diagram showing how the on-context and off-context models differ: on-context models are trained on each non-normal context (spring, wrist, spring+wrist) and tested on the same context. Off-context models are trained on the normal context and then tested on each of the non-normal contexts. (B) Average across non-normal contexts of the Pearson correlation coefficient of the prediction of muscle activity from all numbers of brain synergies up to the total dimension of the brain data. The two black lines represent the baselines for on-context and off-context, when predicting the muscle activity from the full brain data. (C) Same as in B but for the prediction of muscle activity from the full brain data and through muscle synergies (see Methods). (D) Same as in B and C but now showing the prediction of finger velocity from brain synergies. (E) Same as in B, C, and D, but now for the prediction of finger velocity from muscle synergies. (F) Heatmap for the spring+wrist context, like the one shown in Figure 4D, showing the correlation coefficient of the prediction of muscle activity from brain synergies and through muscle synergies, normalized to the off-context baseline prediction. A value over 100 means a better prediction than the baseline off-context model that uses all brain data and predicts muscle activity directly. **Abbreviations**: PCA = principal component analysis; NMF = non-negative matrix factorization; dPCA = demixed PCA; Syn. = synergy.

On-context models were used as a baseline and refer to those trained using the first 80% of the data in each non-normal context and then tested on the last 20% of the data in the same context. Similarly to what we did for the denoising experiment, we trained off-context models to predict finger velocity and muscle activity from the full brain and muscle data (models 1, 2, and 7 in Table 1), which are referred to as the baseline on-context models. Additionally, we trained off-context models to predict finger velocity and muscle activity from brain synergies and through muscle synergies (models 3, 4, 5, 6, and 8 in Table 1; Figure 5A). We measured each model’s performance using the Pearson correlation coefficient between the predictions and the ground truth and compared the performance between on-context and off-context models, as well as between models trained on synergies and models trained on the full data. We tested generalization on one day for Monkey N containing all four task contexts sequentially.

## Results

### Signal Quality Assessment

As an initial step, we sought to verify that the recorded physiological signals were related to the monkey finger movements during the task. The recorded EMG in Monkey N visibly modulated activity with finger movements (Figure 2A). For example, FDP, a flexor muscle targeting middle-ring-small (MRS) flexion, increased activity when moving fingers towards flexion (Figure 2A, top), while EDC, an extensor muscle targeting MRS extension, increased activity when extending MRS (Figure 2A, bottom). Additionally, when using all EMG channels, the muscle activity significantly predicted finger velocity using a ridge regression with history (correlation coefficient CC = 0.60±0.08 across days; p < 1E-4; Figure 2B, right). Finally, we could use the muscle activity to accurately classify the correct kinematic target 97.64%±1.17% of the time (average across n=3 days; chance level 28.2%, nine total targets; Figure 2C).

Similarly, we wanted to verify the intracortical data reflected finger movements too. We recorded intracortical spiking band power (SBP; Nason et al., 2020) from 96 channels in the hand area of the motor cortex for two monkeys: N and W. The “active” channels for each, measured as those that had at least one spike per second on average, are shown in Table 2. With Monkey N, the SBP predicted finger velocities with a slightly lower correlation (CC = 0.53±0.09; p < 1E-4; Figure 2B) compared to the predictions from muscle activity. Monkey N’s SBP could also significantly predict muscle activity (CC=0.71±0.05; p < 1E-19) and accurately classify the task target for each trial (acc = 91.3%±4.6%; Figure 2C). Monkey W’s SBP showed similar predictive capabilities compared to Monkey N, significantly predicting finger velocities (CC = 0.58±0.08; p = 1E-3; Figure 2D) and detecting the correct kinematic target (acc = 86.2%±0.9%; Figure 2C). Overall, these results show that the recorded physiological signals were related to the monkey finger movement during the task.

**Table 2:** Number of active intracortical channels per day and monkey.

|  | Monkey N (total 96) | Monkey W (total 96) |
| --- | --- | --- |
| Day 1 | 53 | 53 |
| Day 2 | 37 | 29 |
| Day 3 | 36 | N/A |

### Relation Between Synergies and Movement

Having shown that the recorded signals were, in fact, related to the monkey finger movements, we were next interested in exploring whether the dimensionality reduction methods could reveal distinct relationships between synergies and movements, particularly in the muscle data. Across all three days and methods (PCA, NMF), the extracted muscle synergies consistently exhibited an explainable structure closely related to the task. Specifically, two synergies were predominantly composed of either flexor or extensor muscles (e.g., synergies 1 and 2 in Figure 3A, respectively), reflecting the flexion-extension nature of the center-out task performed by the monkeys. The third synergy was more broadly distributed among muscles, even those involved in opposing movement patterns, suggesting a supportive role in movement execution. This follows from a more direct relationship between muscles and movement (e.g., FDP is known to be involved in finger flexion) and was quantified by computing the correlations between individual muscle channels and specific movements (Figure 3B). In contrast, the individual brain channels are believed to have a more obscure relationship with movement, which hinders a similar interpretation to the muscle synergies in the context of the task, and was further validated by the lower correlations with movement, compared to the muscle synergies (Figure 3B).

This relationship between brain and movement may be obscure due to an intermediate modulation through brain and muscle synergies. We explored this hypothesis by investigating the relationship between brain and muscles directly, or through synergies. We computed the correlations between all possible pairs of brain and muscle channels (Nx7 pairs for each day, where N corresponds to brain channels in **Table 2**). We computed the number of brain and muscle synergies needed to explain at least 90% of the variance in PCA for each day, and then computed the correlations between all pairs of the activations of these synergies (PxQx3 for each day, where P is the number of brain synergies and Q the number of muscle synergies).

We found generally low (CC <= 0.4; Figure 3C) but significant correlations on most pairs: 93% for brain to EMG, 92% for brain to muscle synergies, 72% for brain synergies to EMG, and 71% for brain synergies to muscle synergies. Interestingly, we also found significantly higher correlations between brain and muscle activity than between all other pairings: higher than brain to muscle synergies, p<1E-20; higher than brain synergies to EMG, p < 1E-20; higher than brain synergies to muscle synergies, p < 1E-20. The stronger relationship between brain and muscles than between using any type of synergy as an intermediate step raises the question as to whether synergies can actually be more useful for decoding than using all the available data, or at least whether they can simply help compress the data more effectively, as has been suggested previously (Cozza et al., 2020; Tang et al., 2021).

### Compression and Denoising Experiments

The ability to compress data effectively is useful by itself, as it can greatly lower power requirements and enable simpler decoding models (Casson et al., 2010; Thies and Alimohammad, 2019). Here, we first compared the different dimensionality reduction methods’ abilities in compressing brain and muscle data in two ways: we first measured the VAF and studied how the total explained variance changed when increasing the number of synergies. Then, we evaluated compressibility end-to-end by measuring the performance of decoders trained on varying numbers of synergies as inputs (Figure 4A). In terms of VAF, three out of seven (∼43%) muscle synergies consistently explained >90% of the variance, across all days and methods (Figure 4C, right). For brain synergies, ∼54% of the total dimensions were needed on average to explain 90% of the variance (Figure 4C, left) for monkey N, while ∼51% were needed for Monkey W (Figure 4E, left). Note that dPCA is extracted from the trial-averaged data and thus cannot explain the full variance of the data, even with all possible synergies. In terms of end-to-end compression ability, we found that all three methods were good at compressing brain data, achieving 95% of the maximum prediction correlation with a relatively small percentage of the total dimensions (Figure 4, B and E), resulting in an average compression ratio of 3.43:1 across monkeys and outputs (Table 3). When compressing muscle data, PCA and NMF had a smaller effect than on the brain data, achieving an average compression ratio across days of 1.75:1 (Table 3). Overall, dimensionality reduction methods were effective in compression, with the brain data showing greater benefits and dPCA showing the best performance.

**Table 3:** Average compression ratios across methods and monkeys.

| Prediction Type | Monkey | PCA | NMF | dPCA |
| --- | --- | --- | --- | --- |
| Brain to Velocity | N | 2.23:1 | 2.23:1 | 4.83:1 |
|  | W | 2.90: 1 | 2.90:1 | 4.14:1 |
| Brain To EMG | N | 2.90:1 | 2.90:1 | 5.80:1 |
| EMG to Velocity | N | 1.75:1 | 1.75:1 | N/A |

Given the promising results in compressing brain and muscle data, we next examined whether dimensionality reduction could also enhance prediction accuracy by reducing noise in the data, i.e., whether it could “denoise” brain and muscle data (De Clercq et al., 2005; Costa-García et al., 2023). Specifically, we were interested in determining if fewer synergies, representing a compressed and potentially less noisy version of the original data, could lead to better predictions compared to using all dimensions. We trained models either using the full dataset or on a reduced set with varying number of synergies (see Methods) and compared their prediction performance. Our results showed that, on both monkeys, none of the dimensionality reduction models outperformed the baseline of using the full data (Figure 4B, D, and E; Supp. Figure 1).

When predicting muscle activity (Figure 4B, top row and heatmap in 4D; Supp. Figure 1) and finger velocity (Figure 4B, bottom row and Figure 4E right), we observed a similar trend: as the number of synergies increased (and thus also the VAF), the prediction performance improved, but never above the baseline (indicated by the black lines on the line plots, 100% in heatmap). This lower performance was consistent across dimensionality reduction methods, with PCA and NMF showing nearly identical behavior and dPCA generally performing better with fewer synergies.

### Generalization Experiment

Previous research suggests that even if synergies cannot help with denoising, they may be stable across tasks (Gallego et al., 2018), which may enable higher generalization across tasks than using all data. We tested this hypothesis by having Monkey N perform the same task under different contexts: normal, with a spring resisting flexion, a 23-degree change in wrist angle, and both wrist and spring changes simultaneously. We trained decoders based on the full data or a set of synergies for both brain and muscles and predicted finger velocities, muscle activity, and muscle activity through muscle synergies. Off-context models were trained on normal trials, while on-context models were trained in the context they were tested on (see Methods, Figure 5A). Overall, we found that, regardless of context, as the number of synergies increased, the correlation with finger velocities and muscle activity increased but did not surpass correlations of the off-context models trained on full data (Figure 5B-F; Supp. Figure 2). Therefore, there does not seem to be any obvious performance enhancing advantages of using synergies to tackle the problem of decoding generalization. The only exception is shown on Figure 5F, where NMF demonstrated a modest improvement in correlation in the spring+wrist context, compared to using the full dataset off-context (1-10%), but this effect was not observed in the other contexts (Supp. Figure 2).

## Discussion

In this study, we tested the ability of three popular dimensionality reduction methods to compress and denoise brain and muscle activity from implanted electrodes, as well as their capacity for generalizing to unseen contexts. Our results demonstrate that while dimensionality reduction methods can effectively compress neural and muscle data, they do not improve decoding accuracy compared to using all available data or help generalize across tasks.

Notably, while PCA and NMF performed comparably in compression, dPCA, a supervised approach, generally showed better performance with fewer synergies, suggesting it may be a preferable approach when compressing for decoding models. Overall, these findings indicate that while synergies provide efficient data compression, they do not help data denoising for better performance than using the full dataset.

Brain and muscle synergies have been suggested in the field of neuroprosthetics as a practical way of solving different issues in performance. For example, (Degenhart et al., 2020) showed that brain synergies can be used to maintain BMI decoder performance across days by transforming each new day’s brain activity to the synergy space. On the muscle side, (Ajiboye and Weir, 2009) showed that muscle synergies can be used to discriminate between postures of the American Sign Language and proposed their use as a control input for prosthetic devices.

Here, we show that even though muscle synergies can be easily interpreted in terms of their kinematic meaning (in our case, two synergies for finger flexion, plus another for extension, Figure 3, A), the correlations between the brain and muscle activations were still always higher than between the brain and muscle synergy activations (Figure 3, C). These higher correlations suggest that a prosthetic control strategy that uses muscle synergies rather than muscle activation for control may be less related to brain activity, which can be undesirable when developing control strategies that are easy and natural to use, especially if the decoding performance with synergies is worse.

Additionally, we showed that synergies extracted using dimensionality reduction techniques were not useful for denoising brain or muscle data from invasive electrodes. Synergies also did not appear to be useful in most cases for generalizing decoders for the same data across contexts. If these techniques could remove unrelated signals (“noise”) from the data, then we would have seen better performance when projecting the data into a lower dimension than when using the full data. This denoising phenomenon has been shown in other studies (Winkler et al., 2011; Damon et al., 2013; Niegowski et al., 2015; Islam et al., 2021; Costa-García et al., 2023), mostly in the context of less invasive recording methods. In our study, however, we found that removing dimensions to the data almost always results in worse performance (Figure 4), which could be driven by the higher inherent signal-to-noise-ratio (SNR) of invasive methods versus their noninvasive counterparts.

Overall, if the dimensionality reduction methods improved the stability of the decoders for brain and muscle across tasks, then we would have seen better generalization performance when using these methods than when using the full data. The stability of the lower dimensional projections of brain data across tasks has been shown in previous studies (Gallego et al., 2018), and others have shown that decoder stability across time can be achieved by reducing the dimensionality and then adjusting future data points. Here, however, we show that dimensionality reduction methods on their own did not help a decoder generalize better than when using the full data in most cases. The results of denoising and generalization collectively suggest that higher-dimensional components, which are typically considered to lack task-related information (Barradas et al., 2020), do contain relevant information for decoding.

Although we were not able to achieve improvements by using synergies for denoising and generalization, these methods were still a great tool for compression. We found that all dimensionality reduction methods achieved good results, showing high compressibility for brain data (3.48:1 for Monkey N, 3.31:1 for Monkey W) and muscle data (1.75:1 for Monkey N; Figure 4). These results suggest that the brain data can be greatly compressed with no loss in decoding performance, which can be very valuable for more complex neural-network-based models. The size of these models is usually dictated by the dimension of the feature vector (Guduru, 2006), and bigger models increase the risk of overfitting to the training data (Srivastava et al., 2014) and require more power to run in real-time (Casson et al., 2010; Gabert et al., 2020). Models that use less power are going to be the key to translating brain-machine interfaces into everyday patient use. Thus, using a linear dimensionality reduction technique as a first layer, potentially even implemented in hardware, that transforms the input features into a lower dimensional space without losing performance can be a key addition to existing complex models. A similar process has been used in other applications to improve model size (Mladenić, 2006).

The three dimensionality reduction methods we tested, PCA, NMF, and dPCA differ in the loss function they minimize and the restrictions they impose: PCA restricts the decoder to be the transpose of the encoder; NMF restricts the activations and the weights to be positive; dPCA allows for different encoder and decoder but computes the transformations subject to a task parameter of interest. In our results, we found dPCA to do the best in terms of compression, denoising, and generalization: it achieved higher compression rates than PCA and NMF (5.32:1 vs 2.57:1 for Monkey N; 4.14:1 vs 2.90:1 for Monkey W) and performed the closest to the full data in the denoising and generalization experiments (Figures 4 and 5). These better results likely follow from the ability of dPCA to have different decoder and encoder transformations, the fact that it is trained in a supervised manner, and the possibility of using task parameters (target locations, in our case) to extract transformations relevant to the task.

This study has some limitations that should be acknowledged. First, we did not test the full range of dimensionality reduction techniques available. We focused on PCA and NMF, which are commonly applied to brain and muscle data, respectively, and included dPCA for brain data to leverage task-specific information. However, other methods, such as more advanced manifold learning techniques or neural network-based approaches (Pandarinath et al., 2018; Abbaspourazad et al., 2023), may offer different or improved insights into data structure and decoding performance, as well as allow for potential non-linear readouts from synergies to behavior. Second, our muscle dataset was limited to a single monkey that had synchronized recordings of muscle activity, brain signals, and kinematics. Although these experiments provide a valuable foundation, the generalizability of our findings is limited by the small sample size. We acknowledge the challenges and costs associated with such experiments, and we aim to expand our study to include additional subjects in future research to strengthen the validity of our conclusions.

In conclusion, the results of this study show that dimensionality reduction methods can effectively compress high-dimensional physiological data for brain-machine interfaces and neuroprosthetics, achieving substantial reductions in data size while retaining predictive power. However, these methods showed limited utility for denoising or improving generalization, as none outperformed the baseline (i.e., using all available data) for predicting EMG or finger velocity. Although synergies approximated full data performance in cross-context tasks, there was no significant advantage in enhancing generalizability. These findings highlight both the value and the limitations of dimensionality reduction for neural data applications, particularly in tasks requiring compact yet informative representations.

## Acknowledgements

We would like to thank Eric Kennedy for animal and experimental support. We thank the University of Michigan Unit for Laboratory Animal Medicine for expert surgical and veterinary care. We thank Samuel Nason-Tomaszewski for his guidance and expertise. LHC was supported by Agencia Nacional de Investigacion y Desarrollo (ANID) of Chile. MMK was supported by NSF NCS. MJM and HT were supported by NSF grant 1926576. MSW was supported by NIH grant T32NS007222. NGK and TAK were supported by NIH grant R01NS105132. PGP and CAC were supported by NSF NCS and NSF BRAID. CK was supported by NSF grant 1804053.

