## Supplemental Figures for "Exploring Synergies in Brain-Machine Interfaces: Compression vs. Performance"

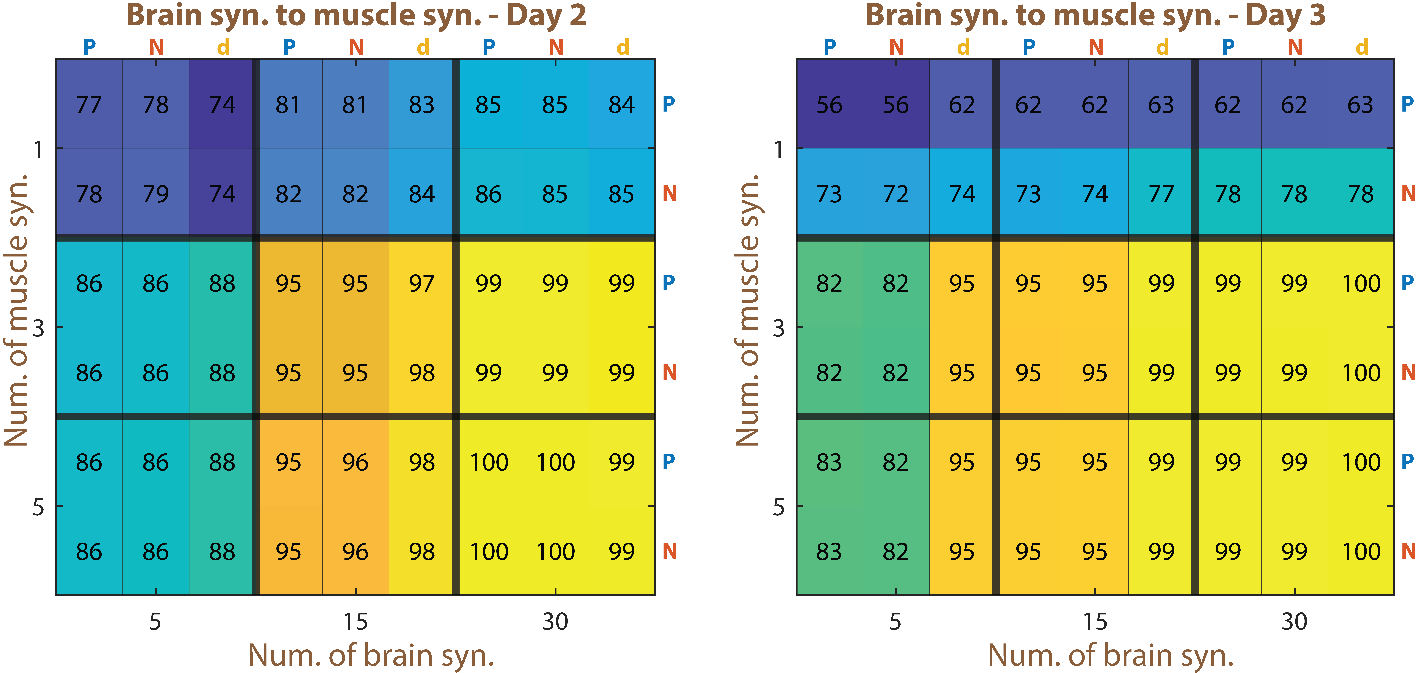

Supp. Figure 1: Heatmap of muscle activity prediction from brain synergies to muscle synergies for day 2 (left) and 3 (right). Each cell value corresponds to the correlation of predicting muscle activity through brain and muscle synergies, normalized to the result of predicting muscle activity directly from brain channels. A value of 100 would mean that it matches the baseline perfectly. P represents PCA, N represents NMF, and d represents dPCA.

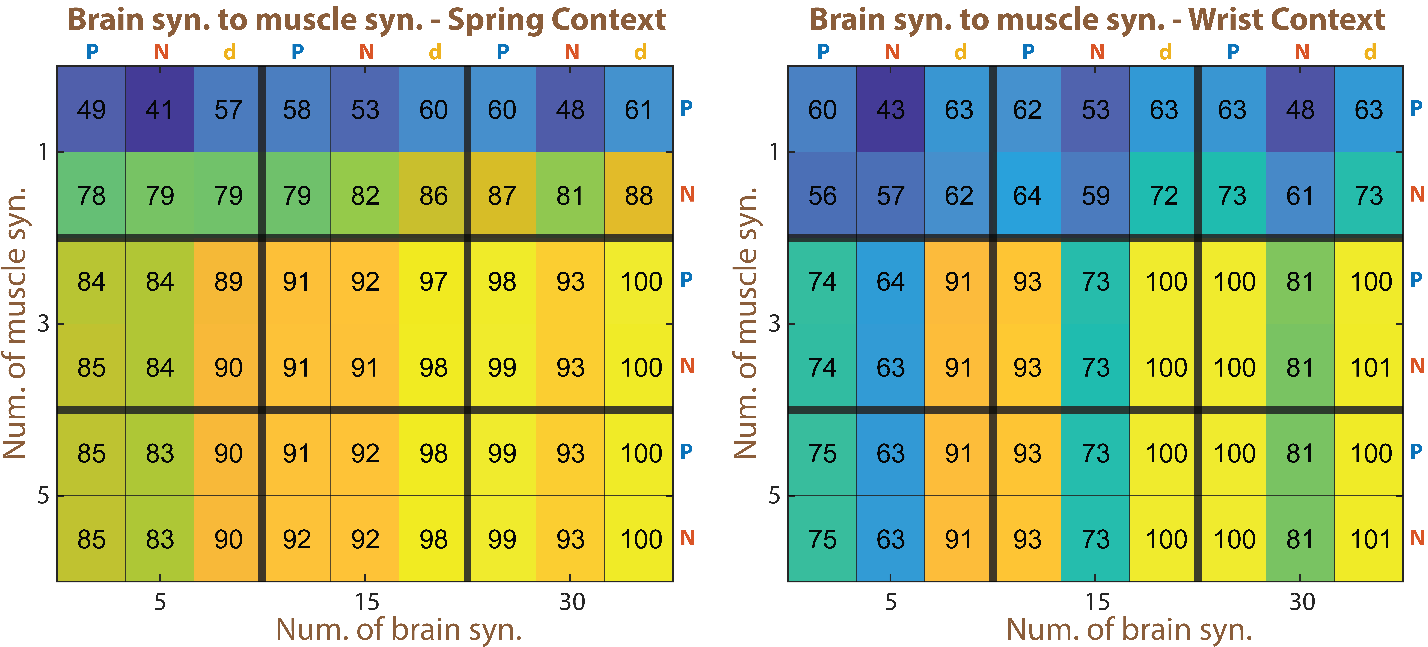

Supp. Figure 2: Heatmap of muscle activity prediction from brain synergies to muscle synergies the spring (left) and wrist (right) contexts on day 1. Each cell value corresponds to the correlation of predicting muscle activity through brain and muscle synergies, normalized to the result of the off-context muscle activity prediction directly from brain channels. A value of 100 would mean that it matches the baseline perfectly. P represents PCA, N represents NMF, and d represents dPCA.
